# NLRP1 is activated by palmitic acid and induced in human metabolic dysfunction-associated steatohepatitis

**DOI:** 10.1101/2024.11.11.622769

**Authors:** Miriam Pinilla, Eulalia Campos Baños, María Antonia Martínez-Sánchez, María Sánchez-Villalobos, Alba Oliva-Bolarín, Sara Rico-Chazarra, Antonio José Ruiz-Alcaraz, Carlos Manuel Martínez, Adriana Mika, María Dolores Frutos, Bruno Ramos-Molina, María Ángeles Núñez-Sánchez, Ana Belén Pérez-Oliva

## Abstract

Metabolic dysfunction-associated steatotic liver disease (MASLD) is the most prevalent cause of liver disease worldwide. This progressive condition ranges from simple steatosis to a more advanced stage, known as metabolic dysfunction-associated steatohepatitis (MASH), which is characterized by inflammation, hepatocellular ballooning, and hepatic steatosis. Several studies have demonstrated the involvement of inflammasomes in MASH development. Recently, the NLRP1 inflammasome has gained attention as an important sensor in various human inflammatory conditions, though its role in metabolic diseases like MASLD remains unclear. In this study, we identified significantly higher mRNA and protein levels of NLRP1 in liver samples from patients with MASH compared to those with normal or steatotic livers. Furthermore, *NLRP1* mRNA levels correlated with hepatic palmitic acid (PA) levels. We also showed that NLRP1 inflammasome expression is mediated by PA in both human HepG2 cells and human liver organoids. Importantly, we found that NLRP1 was activated by PA, but not by other saturated fatty acids like myristic acid, and that PA-induced NLRP1 activation was inhibited by oleic acid. These findings uncover a previously unknown role of the hepatic NLRP1 inflammasome in the human liver. However, further research is needed to fully understand the complex interactions between NLRP1, inflammation, and metabolic processes in the development of MASLD.

## INTRODUCTION

Metabolic dysfunction-associated steatotic liver disease (MASLD) is a leading cause of chronic liver disease worldwide [1]. It represents a progressive disease characterized by lipid accumulation in the liver [2], and it is linked to other metabolic conditions such as obesity, insulin resistance, or dyslipidemia [3]. MASLD ranges from simple steatosis to more severe forms such as metabolic dysfunction-associated steatohepatitis (MASH), which can progress to fibrosis and cirrhosis and, ultimately, to liver hepatocellular carcinoma [4]. Of note, only one FDA-approved pharmacological treatment for MASH with advanced fibrosis is currently available [5]. Therefore, research efforts are needed to identify molecular mechanisms that allow us to develop novel pharmacological treatments for MASH.

Currently, although the physiopathology of MASLD is not yet fully understood, several factors have been described to be involved in the development and progression of the disease, including chronic inflammation and an exacerbated immune response. In this context, it has been suggested that the activation of inflammasomes plays a central role in MASLD, as well as lipotoxicity and insulin resistance activate innate immunity [6]. Inflammasomes are molecular structures localized in the cytosol, which classically have been described as multiprotein complexes mediating immune system activation after infection or tissue injury. Inflammasomes are principally formed by danger signals sensors (NLRs), the adaptor protein ASC, and the effector proteins caspase-1, -4, -5, and -11 [7]. Nucleotide-binding oligomerization domain-like receptor family pyrin domain-containing 1 and 3 (NLRP1 and NLRP3) are intracellular sensors of danger signals from pathogens (pathogen-associated molecular patterns or PAMPs) or tissue damage (damage-associated molecular patterns or DAMPs), and a key component of the innate immune system. Their activation leads to the cleavage of pro-Caspase-1 into its active form, Caspase-1, which then processes pro-inflammatory cytokines such as pro-IL-1β and pro-IL-18 into their active forms. Upon Caspase-1 activation through inflammasomes, cells undergo membrane rupture and lysis, releasing lactate dehydrogenase (LDH).

Several studies have demonstrated that NLRP3, the most studied inflammasome, plays a crucial role in the development and progression of MASLD, as its activation promotes hepatic steatosis and inflammation in various mouse models of MASH [8]. Remarkably, inhibition of NLRP3 with MCC950 has been shown to reduce liver inflammation and reverse fibrosis in a mouse model of MASH [9]. Furthermore, recent studies have shown that patients with MASLD-related fibrosis have significantly higher relative gene expression levels of *IL-1*β and *NLRP3* and propose these inflammatory markers as a potential signal for the early detection of this pathological condition [10]. Moreover, diets rich in saturated fatty acids such as palmitic acid (PA) have been linked to induced NLRP3 [11] [12]. However, to date, other important inflammasomes such as NLRP1 have not yet been associated with this disease.

NLRP1 plays a critical role in initiating and propagating inflammatory responses, but although it was the first sensor described [10], it has been less studied than NLRP3 principally due to the differences in protein structure between humans and mice, which has been used as the main *in vivo* research model [13]. Furthermore, NLRP1 has been linked to metabolic dysregulation, which is a key factor in the development of MASLD although its role is controversial [14] [15]. Other studies have shown that NLRP1 can modulate lipid metabolism, insulin resistance, and glucose homeostasis positively in mice [16], all of which are crucial components in the pathogenesis of MASLD. In addition, administration of an inhibitor of Caspase-1, the effector of the inflammasome, reduces insulin resistance, improves liver fibrosis progression, and delays MASH development in mice in LDLR-/-.Leiden mice fed with a high-fat diet [17].

In this study, we have identified that NLRP1 is highly expressed in hepatic samples from patients with MASH at both the mRNA and protein levels. Additionally, we have found positive correlation exists between hepatic palmitic acid levels and NLRP1, linking palmitic acid accumulation to inflammation in advanced MASLD. Moreover, we have identified that PA induces NLRP1 expression in HepG2 cells and human liver organoids. Notably, PA was able to activate NLRP1, and this activation was prevented by co-treatment with oleic acid (OA). Based on these findings, we propose that NLRP1 plays a critical role in initiating and propagating an inflammatory response driven by PA-induced lipotoxicity in the liver.

## PATIENTS AND METHODS

### Study design and participants

This was a case-control study in 150 patients with obesity undergoing bariatric surgery recruited at the Virgen de la Arrixaca University Hospital (HCUVA, Murcia, Spain) between 2020 and 2023. Details of inclusion and exclusion criteria have been described elsewhere [18]. Briefly, patients between 18-65 years of age with a body mass index ≥ 35 kg/m^2^ or ≥ 30 kg/m^2^ with significant obesity-related comorbidities, scheduled for bariatric surgery who agreed to participate in the study were included in the study. Exclusion criteria included evidence of liver disease other than MASLD, alcohol consumption > 30 g daily for men and > 20 g daily for women, treatment with drugs potentially causing steatosis, or subjects who declined to participate. Patients were divided into three different groups: i) No MASLD (N=53); ii) metabolic dysfunction-associated steatotic liver, MASL (N=43), or iii) MASH (N=54) depending on the histological findings in the liver biopsies according to the Steatosis, Activity, and Fibrosis classification system (SAF Score) [19].

The study was performed in agreement with the Declaration of Helsinki according to local and national laws and was approved by the Ethics and Clinical Research Committees of the Virgen de la Arrixaca University Hospital (ref. number 2020-2-4-HCUVA).

### Anthropometric and biochemical evaluation

Anthropometric and biochemical measurements were performed as previously described [20]. Briefly, anthropometric measurements (weight, height, and waist circumference) were performed by trained staff, and the body mass index (BMI) of the individuals was calculated using Quetelet’s formula (BMI= weight in kg/(height in meters)^2^).

Blood samples were collected on the day of the surgery after an overnight fast of at least 12 hours and then centrifuged to separate the serum. Glucose, total cholesterol (TC), high-density lipoprotein cholesterol (HDLc), triglycerides (TG), aspartate aminotransferase (AST), alanine aminotransferase (ALT), gamma-glutamyltransferase (GGT), alkaline phosphatase (ALP), apolipoprotein A-I (Apo-AI), apolipoprotein B (Apo-B), and albumin levels were performed using the Cobas Analyzer c702 (Roche, Barcelona, Spain) following standardized methods. Non-HDLc was calculated by subtracting HDLc values from TC. Low-density lipoprotein cholesterol (LDLc) levels were calculated using Friedewald’s formula (LDLc = TC - HDLc - TG/5). The levels of glycated hemoglobin (HbA1c) were measured in blood with the glycohemoglobin analyzer HLC®-723G8 (Tosoh Bioscience, Poole, United Kingdom). Insulin levels were measured using the Cobas Analyzer e801 (Roche). The Homeostasis Model Assessment of Insulin Resistance (HOMA-IR) index was used to determine insulin resistance as insulin (μU/mL) x glucose (mmol/L)/22.5 [21].

### Liver biopsies and sample processing

Human liver biopsies were obtained from the patients on the day of the surgery and samples were divided into three parts depending on the downstream analysis. Samples used for histological assessment analysis were fixed in 4% buffered neutral formalin (pH 7.4, VWR, PA, USA), and paraffin embedded. Samples used for RT-qPCR analysis were collected in 1.5 mL nuclease-free tubes containing 1 mL of RNAlater (Sigma-Aldrich, Madrid, Spain), placed at 4 °C overnight and stored at -80 °C until used. Liver samples used for organoid generation were placed in a solution containing DMEM/F-12 + 15 mM HEPES (Gibco, Fisher Scientific, Madrid, Spain) and 10% FBS (Gibco), cut into small pieces, and transferred to a sterile cryogenic vial containing 1 mL of CryoStore CS10 solution (StemCell Technologies, Grenoble, France). The vials were placed in a controlled-rate cell freezing container at -80°C for 24 hours, then transferred to liquid nitrogen for storage until processing.

### Histological assessment and immunohistochemistry

Liver samples embedded in paraffin were sliced into 5 µm sections. For the histological assessment, samples were stained as previously described [20] using hematoxylin and eosin, Masson’s trichrome, Periodic acid–Schiff, Prussian blue, and reticulin staining. The SAF score determination [19] was performed by trained liver pathologists from the HCUVA and the Experimental Pathology Platform of the Biomedical Research Institute of Murcia (IMIB) Pascual Parrilla.

To analyze NLRP1 expression, an indirect immunohistochemistry procedure was carried out in sections from normal, MASL, and MASH liver samples. Briefly, after deparaffination and rehydration, sections underwent a heat-induced demasking antigen procedure by using a commercial solution (Dako Low pH target retrieval solution, Agilent Technologies, Madrid, Spain). After endogenous peroxidase and endogenous background blockage, the sections were overnight incubated with the primary antibody (polyclonal sheep anti-human NLRP1, dilution 1/100; R&D Catalog #: AF6788) at 4 °C, following to incubation with the secondary antibody (polyclonal donkey anti-sheep-HRP labelled, Abcam (Cambridge, UK) dilution 1/100) for 40 min at 37 °C. Immunoreaction was revealed with 3-3’ diaminobenzidine (DAB) and sections were finally counterstained with Harrýs hematoxylin (Epredia In., Madrid, Spain). Positive immunolabelling was identified as a dark-brown precipitated with a cytoplasmic pattern. To establish the degree of NLRP1 expression, the sections were digitized (Pannoramic MIDI II, 3D Histech, Budapest, Hungary) and the whole section was analyzed by using a specialized digital image analysis software (Qpath, ver. 0.5).

### Analysis of fatty acid levels in human liver tissue by gas chromatography-mass spectrometry (GC-MS)

The determination of PA levels in liver samples was performed as previously described [18]. Briefly, total lipids were extracted from liver samples with a 2:1 v/v chloroform-methanol mixture and dried under N_2_ stream. The hydrolysis of extracted lipids was carried out in KOH in methanol [22]. A 10% boron trifluoride in methanol solution was used to obtain FA methyl esters (FAMEs) and gas chromatography-mass spectrometry (GC-MS) analysis of FAMEs was conducted with a GC-MS QP-2010 SE (Shimadzu, Kyoto, Japan).

### Cell culture conditions, treatments, and transfection

Stock solutions of 10 mM PA (Sigma-Aldrich), myristic acid (MA, Cayman Chemical, MI, USA) and OA (Sigma-Aldrich) in 0.1 M NaOH (Sigma-Aldrich) were prepared at 70 °C and stored at -20 °C until use. The stock solution of fatty acid-free bovine serum albumin (BSA, Linus, Cultek, Madrid, Spain), was prepared at 10% by dissolving it in filter-sterilized 0,9% NaCl (Fisher Bioreagents, Madrid, Spain) at 37 °C and stored at - 20 °C until use. On the day of the experiments, PA, MA, or OA were incubated at 70 °C for 10 min to allow the fatty acids to completely dissolve and then were conjugated with 10% BSA in 0.9% NaCl at a 1:1 ratio. The solutions were incubated at 37°C for 1 hour and then filter-sterilized using a 0.22 µm syringe filter (Corning).

HEK293 ASC-GFP cells (Invivogen; kindly provided by Dr. Etienne Meunier, CNRS, Toulouse, France) were maintained in Dulbecco’s Modified Eagle’s Medium (DMEM, Gibco) medium supplemented with 10% fetal bovine serum (FBS, Cytiva HyClone SV30160.03), 1% penicillin-streptomycin (Gibco) and 1% 100X L-glutamine at 37 °C 5% CO_2_. The day before transfection, HEK293 ASC-GFP cells were plated in 24-well or 48-well plates depending on the experiments at a density of 50,000 cells/cm^2^ in culture medium. On the day of transfection, the medium was replaced with OPTIMEM, and cells were transfected with 100 ng of previously described NLRP1 plasmid (pLvB72 hNLRP1) [23] using lipofectamine 2000 according to the manufacturer’s instructions. After approximately 16 hours of transfection, cells were treated or not during 3 hours with albumin-conjugated fatty acids. Albumin controls were obtained by treating cells with 0,1% BSA. In the experiments that used inhibitors, the inhibitors were added one hour before stimulation.

HepG2 cells were obtained from the American Type Culture Collection (ATCC^®^, HB-8065™, American Type Culture Collection, Rockville, MD, USA) and were maintained in DMEM with 25 mM HEPES (Biowest, Nuaillé, France) supplemented with 10% FBS (Gibco), 1% penicillin-streptomycin (Gibco) at 37 °C, 5% CO_2_ and 90% of humidity. Cells were split using trypsin 0.25% EDTA (Sigma-Aldrich) when cultures reached 70-80 % of confluency. HepG2 cells were plated in 6-well, 24-well, or 96-well plates (depending on the downstream analysis) at 50,000 cells/cm^2^ in DMEM complete medium for 24 hours. Cells were then treated for 3 hours with albumin-conjugated fatty acids. Albumin controls were obtained by treating cells with 0.1% BSA. In the experiments that used inhibitors, the inhibitors were added one hour before stimulation. All the analyses were performed in triplicate and in three independent experiments.

### ASC specks

ASC specks formation in HEK293 ASC GFP were monitored after 3 hours of simulations using 4X and 20X objectives using a Leica DMi8 microscope (Leica, Pais). At least 3 different pictures of the nuclei (stained with Hoechst (H3570, Invitrogen)) and specks (GFP) were captured for each condition. The quantification was carried out with the Fiji program. A threshold was applied, and the particle analyzer was used to count the total number of nuclei. The multipoint instrument was used to count the number of specks. Quantification and analysis of ASC specks were performed calculating the number of specks and normalized to the total number of cells stained with Hoechst. Data are shown as fold change of speck-positive cells relative to control.

### Human liver organoid (HLO) isolation, culture and treatment

Cryopreserved human liver samples were thawed at 37 °C and placed in a petri dish containing ice-cold 15 mL of DMEM/F-12 with 15 mM HEPES and 10% FBS. Samples were disassociated using a solution containing DMEM/F-12 with 15 mM HEPES, DNase I (Gibco), and collagenase type IV (Gibco) and incubated at 37 °C for 15 min. Then, samples were resuspended and left for 1 min to allow bigger pieces to precipitate, and the supernatant was transferred to a new tube in ice. This step was repeated 4 times. Once the tissue was completely disaggregated, samples were centrifuged, the supernatant was discarded, and the pellet was resuspended in 10 mL of TrypLE (Gibco) and incubated for 20 min at 37 °C. After this incubation, samples were resuspended and 10 mL of Advanced DMEM with DNase I were added. Samples were centrifuged and the pellet was resuspended in cold Ammonium Chloride Solution (StemCell Technologies) and Advanced DMEM with DNase I and incubated for 5 min in ice. Then samples were centrifuged, the pellets were washed with DMEM/F-12 with 15 mM HEPES and 10% FBS, centrifuged again and resuspended in 2 mL of Advanced DMEM + DNase I. The number of viable cells was determined using Trypan blue solution (Sigma-Aldrich). To initiate the cultures, after centrifuging, the pellet was resuspended in the volume of Matrigel necessary to obtain a density of 20,000 live cells per dome. A total of 40 µL of Matrigel containing cells were placed in each well and the plate was incubated at 37 °C for 30 min before adding 500 µL of HepatiCult Organoid Initiation Media (StemCell Technologies). The medium was changed every 2-3 days until the organoid cultures were ready for passaging (∼ 14 days).

Before performing the experiments, organoid samples were passaged and maintained for at least two passages in HepatiCult Organoid Growth Media (StemCell Tecnologies). Then, organoids were collected and dissociated into small fragments (30 - 100 μm). The number of fragments in suspension was determined using a light microscope. The volume required to obtain 1,000 fragments/well was calculated, transferred to a new tube, centrifuged and the pellet was resuspended in the pertinent amount of Matrigel. For viability assays, 8 µL/well of Matrigel containing fragments were placed in 96-well white opaque plates (Greiner Bio-One, Kremsmünster, Austria). For microscopy analysis and RT-qPCR, 20 µL/well were placed in 48-well plates. The plates were incubated at 37 °C for 30 min before adding HepatiCult Organoid Growth Media and maintained for 5 days changing the media after 2 days. On day 5, the medium was changed to HepatiCult Organoid Differentiation Media (StemCell Technologies) for 10 days before treating the cells with 400 µM or 600 µM of PA. Organoid cultures were incubated for 5 days with the different treatments with change of media every 2-3 days.

### RNA extraction, cDNA preparation, and RT-qPCR analysis

RNA extraction from liver biopsies was performed as described elsewhere [18]. Briefly, liver samples were homogenized in 1 mL of Trizol (Life Technologies, Carlsbad, CA) using an IKA T10 Ultra-Turrax equipment (Janke and Kunkel, IKA Lavortechnick, Germany) and then centrifuged at 12,000 rpm at 4 °C for 5 min. Chloroform was then added to the supernatant and samples were incubated in ice for 5 min before centrifugation at 12,000 rpm at 4 °C for 15 min. The supernatant was collected and transferred to a new tube containing isopropanol, incubating for 10 min at room temperature. Samples were centrifuged at 12,000 rpm for 10 min at 4 °C and the pellet was then washed with 75% ethanol in nuclease free water. After this, samples were centrifuges at 7,500 rpm for 5 min at 4 °C and the pellet was allowed to dry at room temperature for 10 min before resuspending in nuclease free water.

To extract RNA from HLO, organoids were first isolated from Matrigel using ice-cold Cell Recovery Solution (Corning). The solution was added directly into the wells and the plates were placed in ice for 30 min under agitation. After this, samples were collected in 1.5 mL nuclease free tubes and centrifuged at 2,000 x g for 5 min at 4 °C. Samples were then washed 4 times with ice-cold PBS. RNA extraction was performed using the PicoPure RNA Isolation Kit (Applied Biosystem, Waltham, MA, USA) following manufacturer’s instructions. Total RNA from HepG2 cells was extracted using the GenElute Mammalian Total RNA Miniprep kit (Sigma-Aldrich) following the manufacturer’s instructions.

RNA was eluted in RNAse-free water and checked for concentration and purity using the Nanodrop spectrophotometer system (ND-100 3.3. Nanodrop Technologies, USA). Only samples with a ratio of Abs260/Abs280 between 1.8 and 2.1 were used for gene expression assays. Reverse transcription to cDNA was performed using the High-Capacity RNA-to-cDNA kit (Applied Biosystems) following the manufacturer’s instructions. Real-time quantitative polymerase chain reaction (RT-qPCR) amplification was carried out with the Power SYBR Green Master mix (Applied Biosystems) using a Fast 7500 Real-Time instrument (Applied Biosystems) as previously described [24]. The sequences of the primers used in this study can be found in **Table S1.**

### Cell death assay

Cell lysis was evaluated by the quantification of LDH release into the cell supernatant, employing the cytotoxic detection kit (LDH) (11 644 793 001, Roche). HepG2 cells were seeded in 6-well plates at a density of 5 × 10[ cells per well in complete DMEM. The following day, cells were treated with different treatments described in each figure. After 24 hours, 50 µL of cell supernatant was collected and mixed with an equal volume of LDH substrate in a 96-well plate. The mixture was incubated for 10 minutes at room temperature and protected from light. The enzymatic reaction was stopped by adding 50 µL of stop solution. Maximal cell death was determined using whole cell lysates from unstimulated cells incubated with 1% Triton X-100.

### Statistical analysis

Data from the clinical trial were analyzed using the SPSS Statistics 28.0.1.1 software (IBM Corp., Armonk, NY, USA). Data from the *in vitro* studies were analyzed using the Prism 9.0 program (GraphPad Software Inc., Boston, MA, USA). Normality of the study variables was assessed using the Shapiro-Wilk test. The analysis of categorical data was performed by the χ2 test for intragroup differences and the Pearson χ2 test for the intergroup differences. Differences between quantitative variables were assessed using the Kruskal–Wallis one-way ANOVA on ranks in those variables without a normal distribution, whereas ANOVA was used in those parameters following a normal distribution, followed by Bonferroni post hoc analysis for the intergroup differences. Correlations between pairs of variables were carried out using the Spearman correlation test. All values are given as mean[±[standard deviation (SD). A level of p<[0.05 was considered statistically significant. Figures were represented using the Prism 9.0 program (GraphPad Software Inc.).

## RESULTS

### NLRP1 expression is increased in liver tissue samples from patients with MASH and correlates with hepatic palmitic acid concentrations

**Figure 1A** represents the characteristics and progression of MASLD (normal liver, MASL, and MASH). The clinical, anthropometric, and biochemical data of the study patients are shown in **Table S2**. The histopathological findings of these patients are included in **Table S3.** In order to evaluate the alterations in the inflammatory state in different stages of MASLD (53 without MASLD, 43 with MASL, and 54 with MASH), we evaluated the expression of *NLRP1* and *NLRP3* inflammasomes as well as of genes implicated in the inflammatory response in liver samples by RT-qPCR. Our results showed that, while there were no differences in *TNF-a*, *IL-6,* and *NLRP3* expression between the study groups (**Figure S1)**, the expression of *IL-1*β and *NLRP1* was significantly elevated in patients with MASH compared to those without MASLD (**Figure 1B**). Moreover, we observed statistically significant differences in *NLRP1* expression between patients with MASL and patients with MASH (**Figure 1B**).

**Figure 1.**
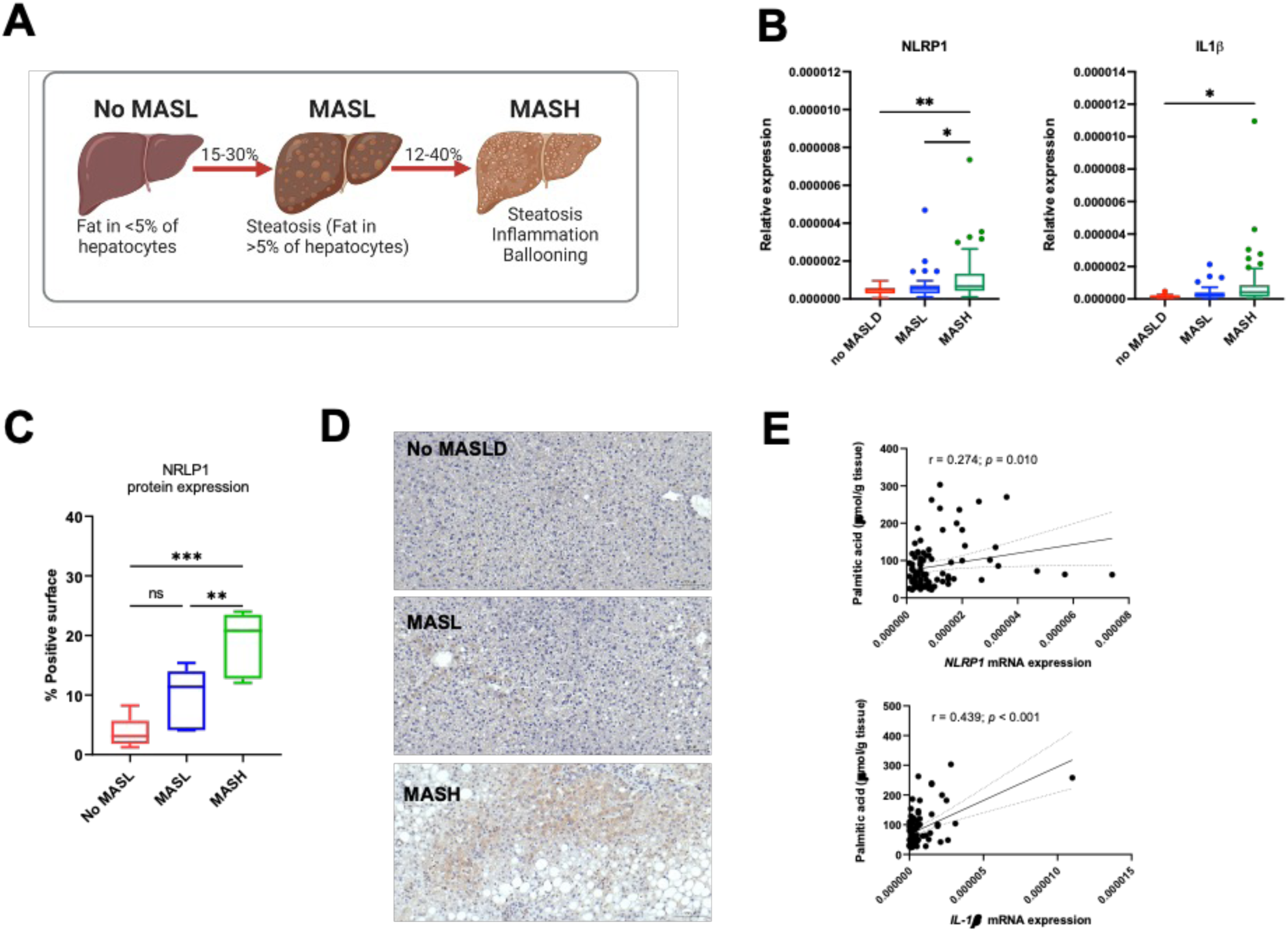
Increased NLRP1 and IL-1β expression in liver samples from MASH patients. **(A)** Schematic illustration of the nomenclature and stages of the progressive form of metabolic dysfunction-associated steatotic liver disease (MASLD). **(B)** Relative expression levels of *NLRP1* and *IL-1*β normalized to 18S rRNA in liver biopsy samples from different patient groups: no MASLD (n=29), MASL (n=44), and MASH (n=54). Data are represented as box and whisker plots using Tukey’s method, where the boxes show the interquartile range (IQR), and the whiskers extend to 1.5 times the IQR. Outliers are represented as individual points. Statistical analysis was performed using one-way ANOVA. p<0.05, p<0.01, p<0.001. **(C)** Quantification of NRLP1 protein expression levels analyzed by immunohistochemistry (no MASLD, n=5; MASL, n=6; MASH, n=6). Quantitative data are presented as box and whisker plots using Tukey’s method, where the boxes represent the interquartile range (IQR), and the whiskers extend to 1.5 times the IQR. Outliers are represented as individual points. **(D)** Representative immunohistochemistry images stained with hematoxylin and NLRP1 (brown) in liver biopsy samples from each study group. Statistical analysis was performed using one-way ANOVA. p<0.05, p<0.01, p<0.001. **(E)** Correlation analysis of palmitic acid (16:0) concentration in hepatic tissue with the mRNA expression levels of *NLRP1* and *IL-1*β, normalized to 18S rRNA. Correlation coefficients (r) are represented as Spearman’s rho, and statistical significance is indicated as follows: *p* < 0.05 (significant), *p* < 0.01 (highly significant), *p* < 0.001 (very highly significant).

To evaluate whether the observed increase in *NLRP1* mRNA expression correlated with protein expression, we determined the levels of NLRP1 by immunohistochemistry staining in a subgroup of samples (No MASLD = 5, MASL = 6 and MASH = 6) (**Figure 1C-D**). On the one hand, our results showed a slight yet not significant increase in the levels of NLRP1 in patients with MASL compared to patients without MASLD (**Figure 1C**). On the other hand, the analysis revealed that, in agreement with the results obtained from mRNA expression analysis, the concentrations of NLRP1 in hepatocytes were increased in patients with MASH compared to those with MASL or without MASLD. Moreover, the quantification of the positive surface area showed an increase of about 15% and 9% in patients with MASH compared to patients without MASLD (*p* < 0.001) and MASL (*p* = 0.009), respectively (**Figure 1C-D**).

One of the hallmarks of MASLD is the intracellular accumulation of lipids in hepatocytes. Particularly, the accumulation of PA has been described to promote lipotoxicity and might be related to the development of MASLD [25]. Thus, we next evaluated if increased expression of NLRP1 and inflammatory genes correlated with hepatic PA levels. Our results showed that, there was a strong positive correlation between the expression levels of *IL-1*β and *NLRP1* mRNA expression and PA levels in the liver (*p* < 0.001 and *p* = 0.010, respectively) (**Figure 1E**).

### Palmitic acid induces NLRP1 expression *in* in vitro models of hepatocytes and human liver organoids derived from MASLD patients

To further investigate the specific effects of PA accumulation on *NLRP1* expression in hepatocytes, we exposed human HepG2 cells and a model of human liver organoids (HLO) to different concentrations of PA (**Figure 2A**). Our results revealed that the exposure of HepG2 to PA significantly increased the expression of *NLRP1* mRNA after 24 hours compared to untreated cells (*p* = 0.036, **Figure 2B**). On the other hand, the analysis in HLO exposed to 400 µM and 600 µM PA showed a slight dose-dependent increase in *NLRP1* expression after 48 hours, although this increase was not significant in any of the concentrations tested (**Figure 2B**), despite the alterations observed in the morphology of differentiated HLO at the highest concentration of PA (**Figure 2C**).

**Figure 2.**
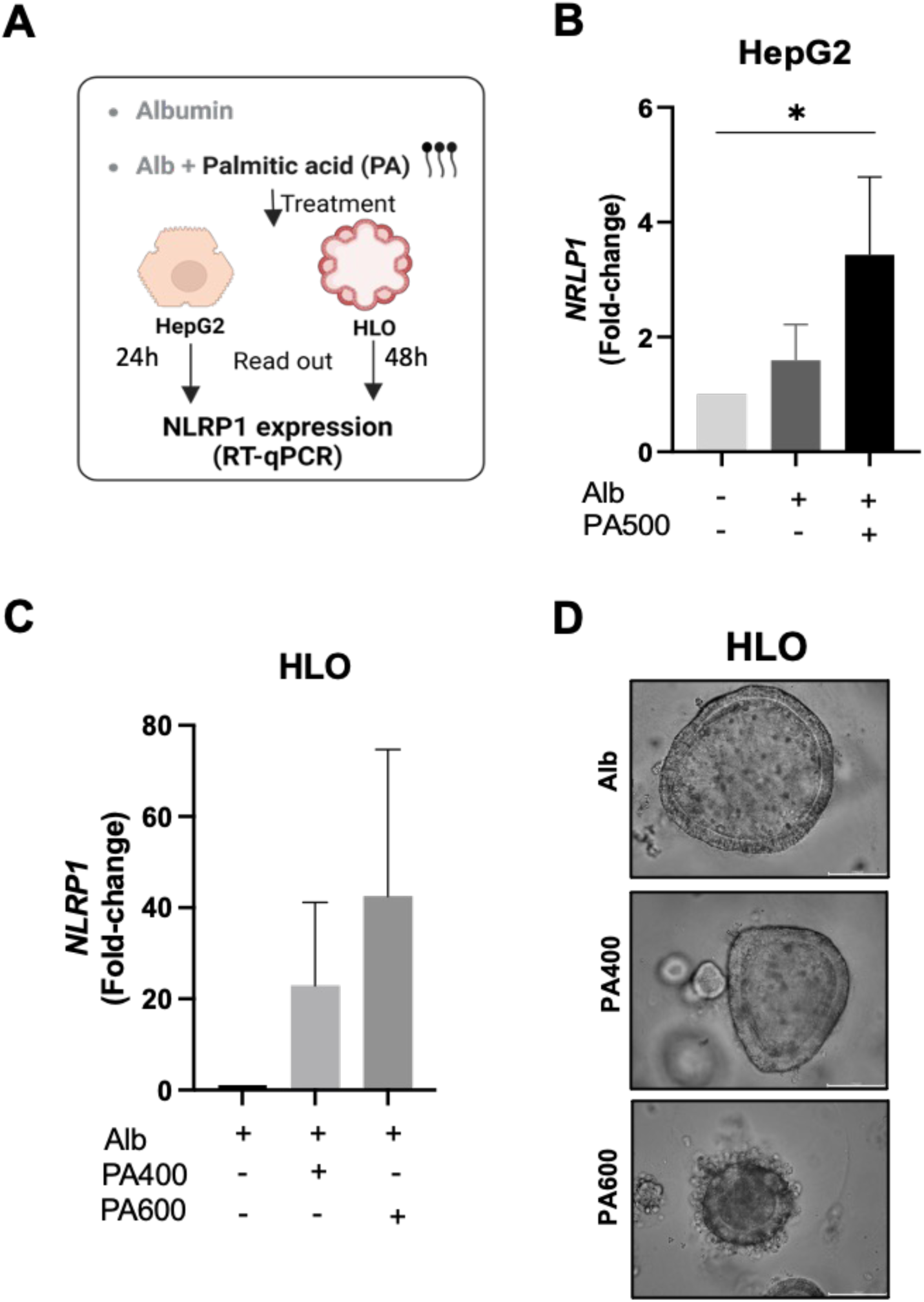
*NLRP1* expression is induced by palmitic acid (PA) in HepG2 cell line and human liver organoids. **(A)** Schematic protocol illustrating the treatment of HepG2 cells or human liver organoids (HLO) with albumin (Alb) or palmitic acid (PA) for 24 or 48 hours. Following the treatment period, *NLRP1* expression is measured by RT-qPCR as the experimental readout. **(B)** Relative gene expression of *NLRP1* normalized to negative control in HepG2 cells treated during 24 hours with Alb or PA (500 μM). **(C)** Relative expression of *NLRP1* gene normalized to albumin in HLO treated with albumin (alb), PA (400-600 μM) during 48h. (D) Representative images of HLOs treated with either Alb or PA (400 or 600 μM) for 48h (bar = 100 μm).

### NLRP1 is activated by palmitic acid, but not other saturated fatty acids such as myristic acid

Considering PA induction on *NLRP1* expression in both HepG2 cells and HLO, we analyzed if MA, another saturated fatty acid, which is linked to MASH although is not lipotoxic [26], could have similar effects. Therefore, we exposed the HepG2 cells to either PA or MA using albumin as a control as indicated in **Figure 3A**. Our results show that PA, but not MA, was able to induce release of LDH, and marker of cell death, in this cell line (**Figure 3B**). In addition, we assessed if PA was also able to regulate ASC oligomerization and specks formation mediated by NLRP1, which will be indicative of NLRP1 inflammasome activation. In order to do that, HEK293 cells expressing ASC-GFP and transfected with human NLRP1 were incubated with either PA, MA or vehicle (albumin), as reflected in **Figure 3C**. We observed that cells treated with PA were able to increase ASC specks formation depending on NLRP1 expression, whereas albumin or MA had no effect (**Figure 3D**). A representative image of each condition measured in **Figure 3D** is shown in **Figure 3E**. **Figure S2** shows a representative image of all the controls of the experiments including cells without NLRP1 expression and treated with albumin, PA or MA.

**Figure 3.**
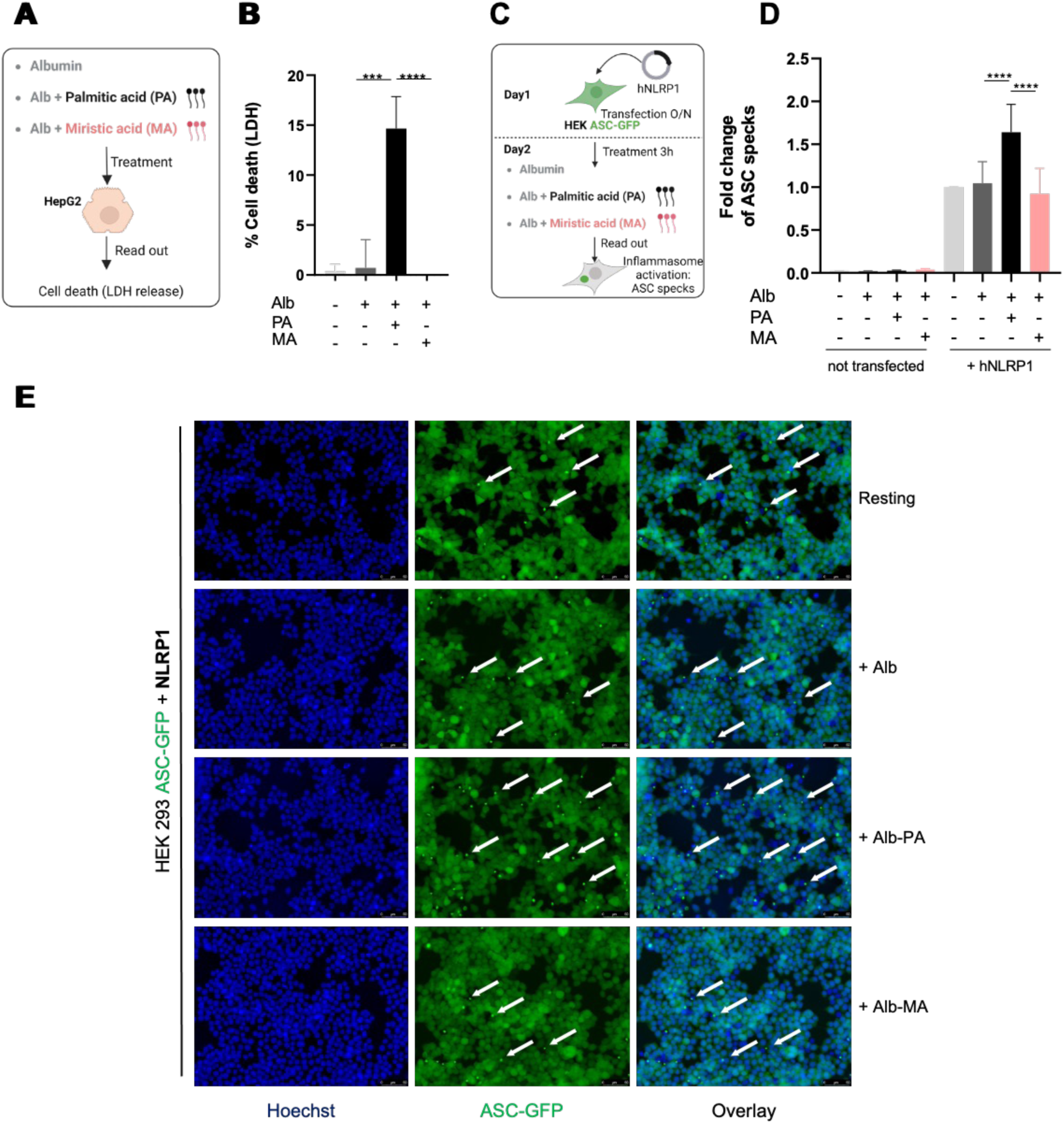
Palmitic acid triggers cell death in HepG2 cells and NLRP1 activation in HEK ASC-GFP cells. **(A)** Schematic representation of the protocol used to treat HepG2 cells: HepG2 cells were treated for 24 hours with either palmitic acid (500 μM) or myristic acid (500 μM), both previously conjugated with albumin (Alb-PA and Alb-MA, respectively). The control group was treated with albumin alone (Alb). **(B)** Measure of cell lysis (LDH release) in HepG2 cells treated for 24h with Alb, Alb-PA (500 μM) and Alb-MA (500 μM). Graphs represent means ± SD from four independent experiments. *P<0.05; **P<0.01; ***P<0.001, ****P<0.0001 by One Way Anova test. **(C)** Schematic representation of the experimental protocol for treating HEK ASC-GFP cells. **(E)** Fluorescence microscopy images and associated quantifications **(D)** of ASC-GFP speck formation in HEK ASC-GFP reporter cells transfected with NLRP1 and treated for 3 hours Alb, Alb-PA (500 μM) or Alb-MA (500 μM). Speck formation is indicated with a white arrow. Images shown are from one experiment and are representative of n=3 independent experiments. At least 3 fields per experiment were analyzed. The number of specks was quantified by counting the green fluorescent dots (ASC-GFP specks) and normalized to the total number of cells stained with Hoechst. Data are shown as fold change of speck-positive cells relative to control (mean ± SD, n=3; *P<0.05, **P<0.01, ***P<0.001, ****P<0.0001, two-way ANOVA).

### Oleic acid prevents NLRP1 activation by palmitic acid

Finally, given that OA has been reported to prevent PA-mediated NLRP3 activation [27], we next assessed whether OA could also prevent NLRP1 activation after PA exposure. For that purpose, we determined cell death by measuring LDH release after treating the cells with PA alone or in combination with OA, as indicated in **Figure 4A**. The results indicate OA abolished PA-mediated cell death (**Figure 4B**). Furthermore, to assess the impact of OA on NLRP1 activation mediated by PA, we treated HEK293 cells expressing ASC-GFP as described in **Figure 4C**. Quantification of ASC specks formation demonstrated that treatment with OA prevents NLRP1 activation induced by PA (**Figure 4D**). A representative image of each treatment condition is reflected in **Figure 4E**. **Figure S3** shows a representative image of all the controls of the experiments including all cells without NLRP1 expression and treated with albumin, PA or OA.

**Figure 4.**
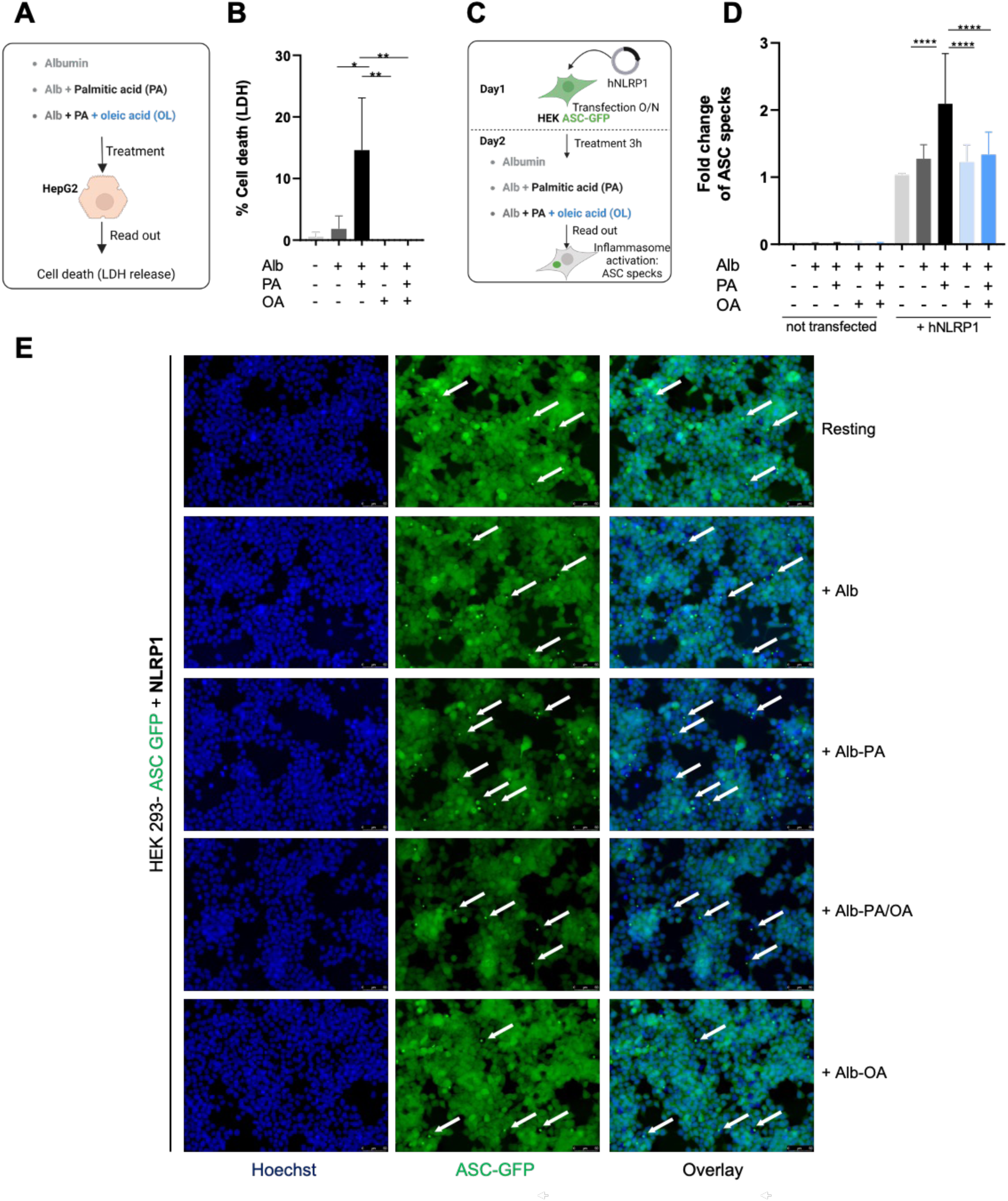
Oleic acid prevents palmitic acid-induced NLRP1 activation in HEK ASC-GFP cells. **(A)** Schematic representation of the protocol used to treat the HepG2 cell line: HepG2 cells were treated for 24 hours with albumin-palmitic acid (500 μM) (alb-PA), oleic acid (250 μM) (alb-OA), or a combination of palmitic and oleic acid (alb-PA/OA). The control group was treated with albumin alone (Alb). **(B)** Measure of cell lysis (LDH release) in HepG2 cell line treated during 24h as described above. Graphs represent means ± SD from four independent experiments. *P<0.05; **P<0.01; ***P<0.001, ****P<0.0001 by One Way Anova test. **(C)** Schematic representation of the experimental protocol for treating HEK ASC-GFP cells. **(E)** Fluorescence microscopy images and associated quantifications **(D)** of ASC-GFP in HEK ASC-GFP reporter cells transfected with NLRP1 and treated for 3 hours with alb-PA (500 μM), alb-OA (250 μM) or a combination of both (alb-PA/OA). Speck formation is indicated with a white arrow. Images shown are from one experiment and are representative of n=4 independent experiments. At least 3 fields per experiment were analyzed. The number of specks was quantified by counting the green fluorescent dots (ASC-GFP specks) and normalized to the total number of cells stained with Hoechst. Data are shown as fold change of speck-positive cells relative to control (mean ± SD, n=3; *P<0.05, **P<0.01, ***P<0.001, ****P<0.0001, two-way ANOVA).

## DISCUSSION

In this study, we have shown for the first time that NLRP1 expression is increased in hepatic samples of patients with MASH at both mRNA and protein levels. In addition, we found that the mRNA expression levels of different inflammatory markers, including *NLRP1*, positively correlated to the concentration of PA in human liver tissue samples. The *in vitro* assays using human HepG2 cells and HLO demonstrated that exposure to PA was associated with an increased expression of *NLRP1*. The use of the stable cell line HEK293 expressing ASC-GFP and transfected with human NLRP1 plasmid showed an increase in ASC oligomerization after PA treatment but no other saturated fatty acids such as MA. Furthermore, the addition of OA was able to inhibit the speck formation in PA-exposed cells.

To the best of our knowledge, this is the first study demonstrating the overexpression of NLRP1 in hepatic samples from patients with MASLD. As inflammation is a key factor in the progression of MASLD, the study of inflammasomes in this disease has gained increased attention over the past years [28]. In this regard, overexpression of NLRP3 as well as related inflammatory markers such as IL-1β have been described to be overexpressed in liver samples from patients with MASLD [29] [30] [31] [32] [33] as well as in animal models [9]. NLRP3 inflammasome activation leads to the generation of proinflammatory cytokines and increased production of reactive oxygen species (ROS), which, in turn, enhance the expression of NLRP3, exacerbating the inflammatory response and promoting MASLD occurrence and severity [34]. However, while most of these studies have been focused on the study of NLRP3, little is known regarding the potential role of NLRP1 in MASLD pathogenesis. Here we have analyzed the mRNA expression of pro-inflammatory cytokines as well as *NLRP3* and *NLRP1* in hepatic samples from obese patients and different degrees of MASLD. On the one hand, our results showed that there was an increase in *NLRP3*, although not significant, independently of the severity of MASLD. This contrasts with previous studies where NLRP3 activation has been related to MASH and fibrosis rather than simple steatosis [33] [30] [31]. This could be explained because some of these analyses were performed at the protein level and mRNA expression might be occurring at earlier stages of the disease [31]. In addition, some of the studies were performed in liver samples from patients with MASH compared to healthy controls, and thus disregarding NLRP3 expression at less severe stages of MASLD [30]. On the other hand, *IL-1*β and *NLRP1* mRNA expression was increased in patients with MASH compared to those patients without MASLD. Furthermore, this elevation in *NLRP1* expression seems to be specific to more advanced stages of MASLD, as differences were also observed between patients with MASH and those with simple steatosis at translational and transductional levels. This is of interest because, while both NLRP1 and NLRP3 lead to the activation of similar downstream pathways, such as the maturation and secretion of IL-1β and IL-18 [35], these results suggest that there may be differences in their regulation and their contribution to MASLD pathology, indicating that NLRP1 might be more relevant in the transition from simple steatosis to MASH than other inflammasomes.

One of the hallmarks of MASLD is the accumulation of free fatty acids in hepatocytes, which are known to be major players in the pathogenesis of the disease [20] [9]. Specifically, the increase in PA concentrations in hepatocytes is largely known for its lipotoxic effect [36]. In this regard, we have previously demonstrated in a cohort of patients with obesity that the concentrations of hepatic PA are higher in patients with MASH. Furthermore, such an increase in PA concentrations was correlated with parameters involved in the progression and severity of MASLD, including lobular inflammation and fibrosis [20]. In this study, we found a strong positive correlation between concentrations of hepatic PA and mRNA levels of both *NLRP1* and *IL-1*β. Furthermore, the use of *in vitro* models of an established line of hepatocytes and HLO from patients corroborates that exposure to PA can induce the expression of *NLRP1* in these models. In this regard, previous research has described the role of PA on NLRP3 activation [30] [11]. However, less is known about the influence of PA on NLRP1 induction and activation. In this regard, the accumulation of PA in hepatocytes has been described to promote mitochondrial dysfunction together with the generation of reactive oxygen species (ROS) [9] and ER stress [37] [38]. Keeping this in mind and considering that specks formation assay has been classically used to evaluate inflammasome activation [39], we explored and demonstrated that PA is able to increase ASC oligomerization dependent on NLRP1 in HEK293 cells. Similar assays were performed using MA without any impact on speck formation. These results could be explained by considering the lipotoxicity of PA compared to MA [26]. Finally, we confirmed that ASC oligomerization and specks formation dependent on NLRP1 by PA were abolished in the presence of OA, something similar to what has been described for NLRP3 [27].

Altogether, our data demonstrates a new role of NLRP1 in MASLD, which is elevated both at mRNA and protein levels and correlates with the severity of the disease and the concentrations of PA in human liver samples. In addition, PA exposure induced NLRP1 expression in HepG2 cells and HLO. Finally, we demonstrated that ASC oligomerization dependent on NLRP1 is induced by PA but not MA and that OA provides protection against such activation.

## Acknowledgments

The authors would like to express their gratitude to all the participants who agreed to participate in this study, as well as the staff at the Virgen de la Arrixaca University Hospital of Murcia (surgeons, endocrinologists, nurses, and clinical biochemists) who have generously participated in the study. We are also grateful for the collaboration of the Biobank Network of the Region of Murcia, BIOBANC-MUR, registered on “Registro Nacional de Biobancos” with registration number B.0000859. BIOBANC-MUR was supported by the “Instituto de Salud Carlos III (proyecto PT20/00109)”, by “Instituto Murciano de Investigación Biosanitaria, IMIB” and “Consejería de Salud de la Comunidad Autónoma de la Región de Murcia”. We also thank Dr. Meunier for kindly providing essential reagents (HEK293 cells expressing ASC-GFP and NRLP1 plasmid).

## Funding

This work was supported by the Spanish Ministry of Science, Project PID2021-126751-NA-100 funded by MCIN/AEI/10.13039/501100011033/ and by ERDF “A way to make Europe” and by the Institute of Health Carlos III (ISCIII) (PI23/00171). A.B.P.-O., B.R.-M., and M.A.N.-S. are supported by the “Miguel Servet” program (CP20/00028, CP19/00098, and CP23/00051, respectively, ISCIII, Spain; co-funded by the Fondo Europeo de Desarrollo Regional-FEDER). M.A.M-S is supported by a PFIS predoctoral fellowship from the ISCIII (FI21/00003, ISCIII, Spain; co-funded by the Fondo Europeo de Desarrollo Regional-FEDER). M.S.V is supported by Rio Hortega project (CM22/00116 ISCIII, Spain; co-funded by the Fondo Europeo de Desarrollo Regional-FEDER). The funding organization played no role in the design of the study, review, and interpretation of the data, or final approval of the manuscript.

## Author contributions

MPM, ECB, AOB, SRC, CMM and AM conceived and carried out experiments; MPM, ECB, MSV, AOB, SRC, AJRA, CMM, AM, BRM, MANS and ABPO conceived experiments and analyzed data. MAMS, AOB, MDF and BRM collected patient data and samples. BRM, MANS, and ABPO were responsible for supervision, project administration, and funding acquisition. All authors were involved in writing the paper and had final approval of the submitted and published versions.

## Conflict of interest

The authors declare that they have no conflict of interest.

## Notes

### Competing Interest Statement

The authors have declared no competing interest.

