## Supplementary figures for "NLRP1 is activated by palmitic acid and induced in human metabolic dysfunction-associated steatohepatitis"

### Slide 1
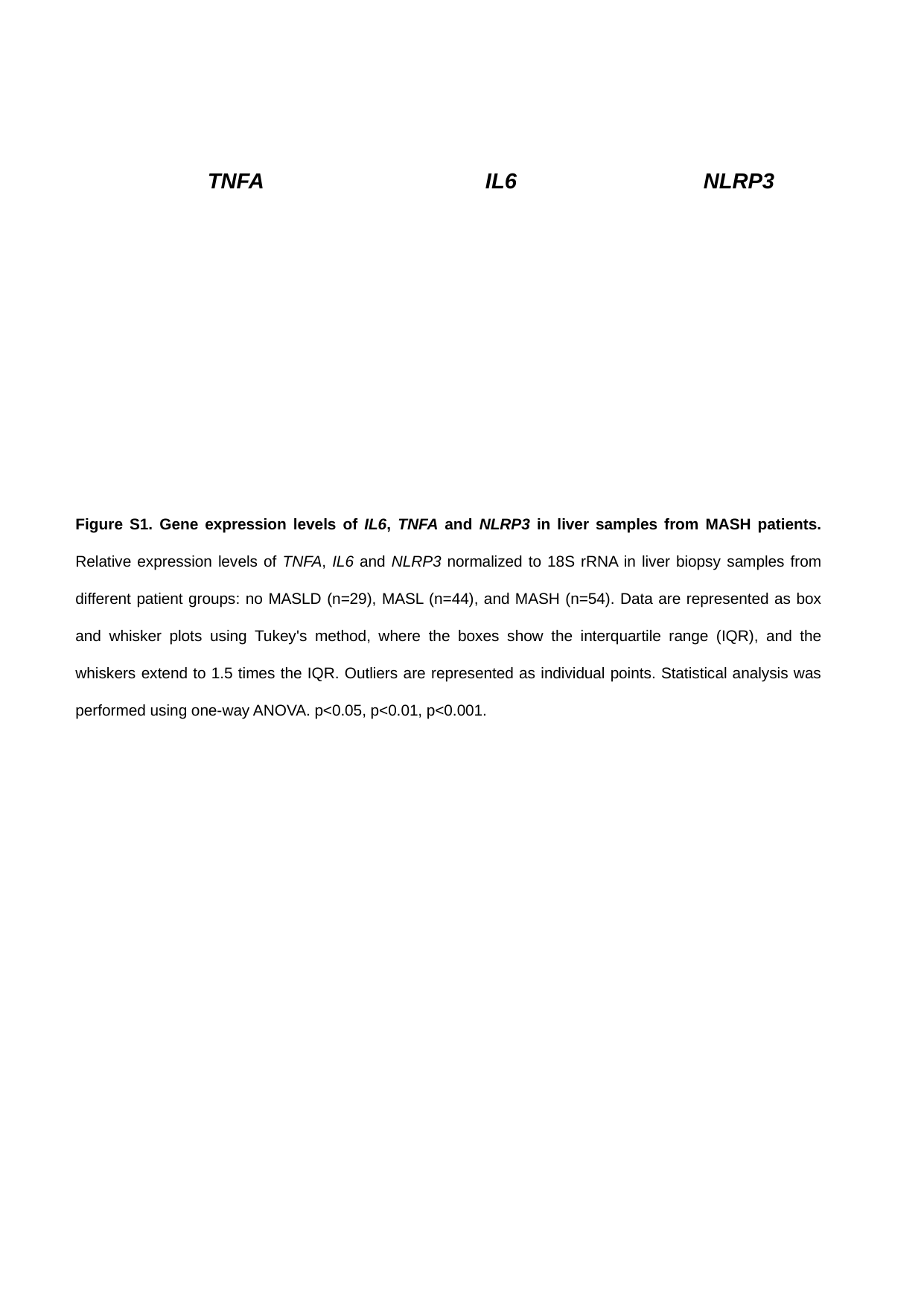

NLRP3
TNFA
IL6
Figure S1. Gene expression levels of IL6, TNFA and NLRP3 in liver samples from MASH patients. Relative expression levels of TNFA, IL6 and NLRP3 normalized to 18S rRNA in liver biopsy samples from different patient groups: no MASLD (n=29), MASL (n=44), and MASH (n=54). Data are represented as box and whisker plots using Tukey's method, where the boxes show the interquartile range (IQR), and the whiskers extend to 1.5 times the IQR. Outliers are represented as individual points. Statistical analysis was performed using one-way ANOVA. p<0.05, p<0.01, p<0.001.

### Slide 2
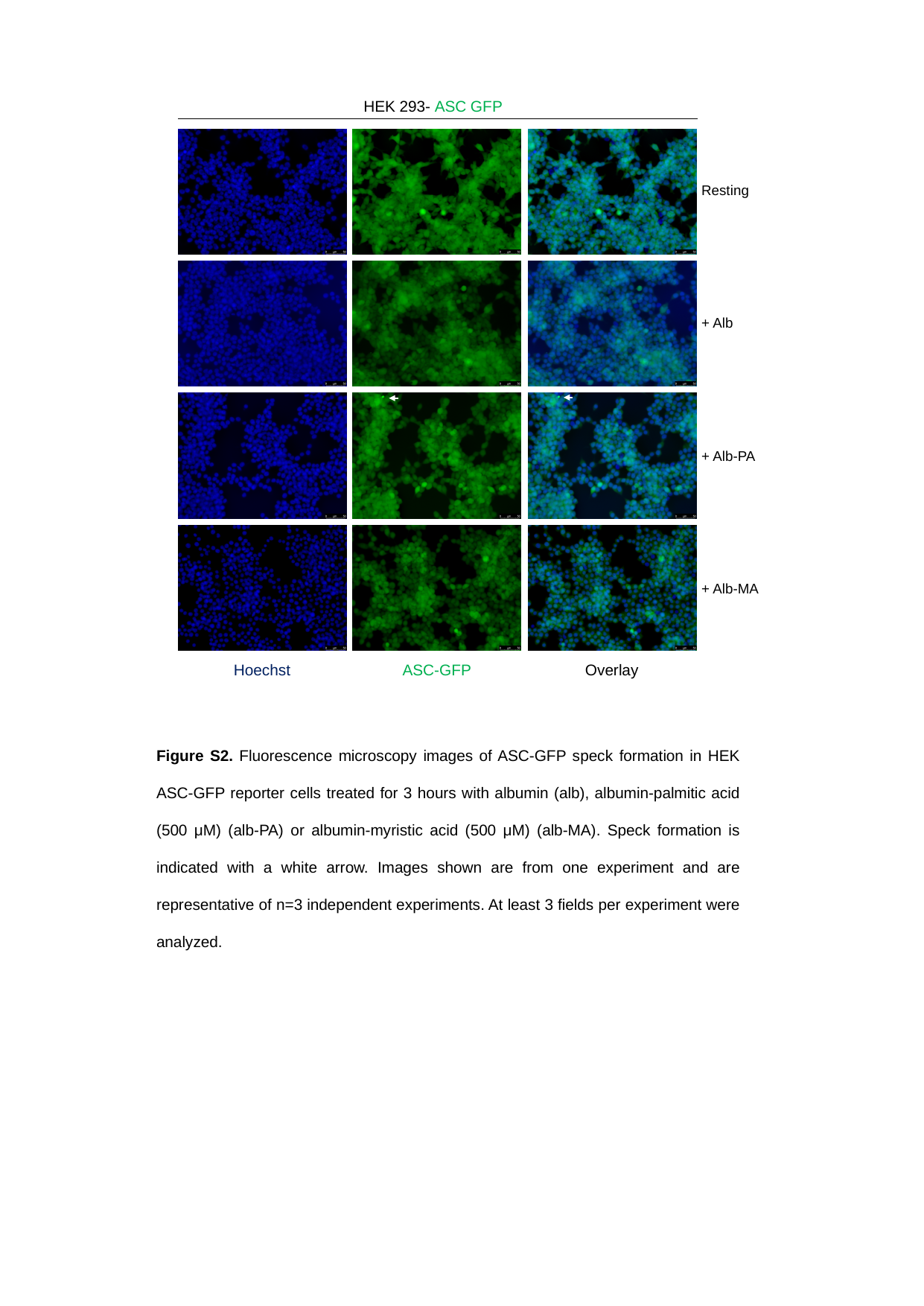

HEK 293- ASC GFP
Resting
+ Alb
+ Alb-PA
+ Alb-MA
ASC-GFP
Overlay
Hoechst
Figure S2. Fluorescence microscopy images of ASC-GFP speck formation in HEK ASC-GFP reporter cells treated for 3 hours with albumin (alb), albumin-palmitic acid (500 μM) (alb-PA) or albumin-myristic acid (500 μM) (alb-MA). Speck formation is indicated with a white arrow. Images shown are from one experiment and are representative of n=3 independent experiments. At least 3 fields per experiment were analyzed.

### Slide 3
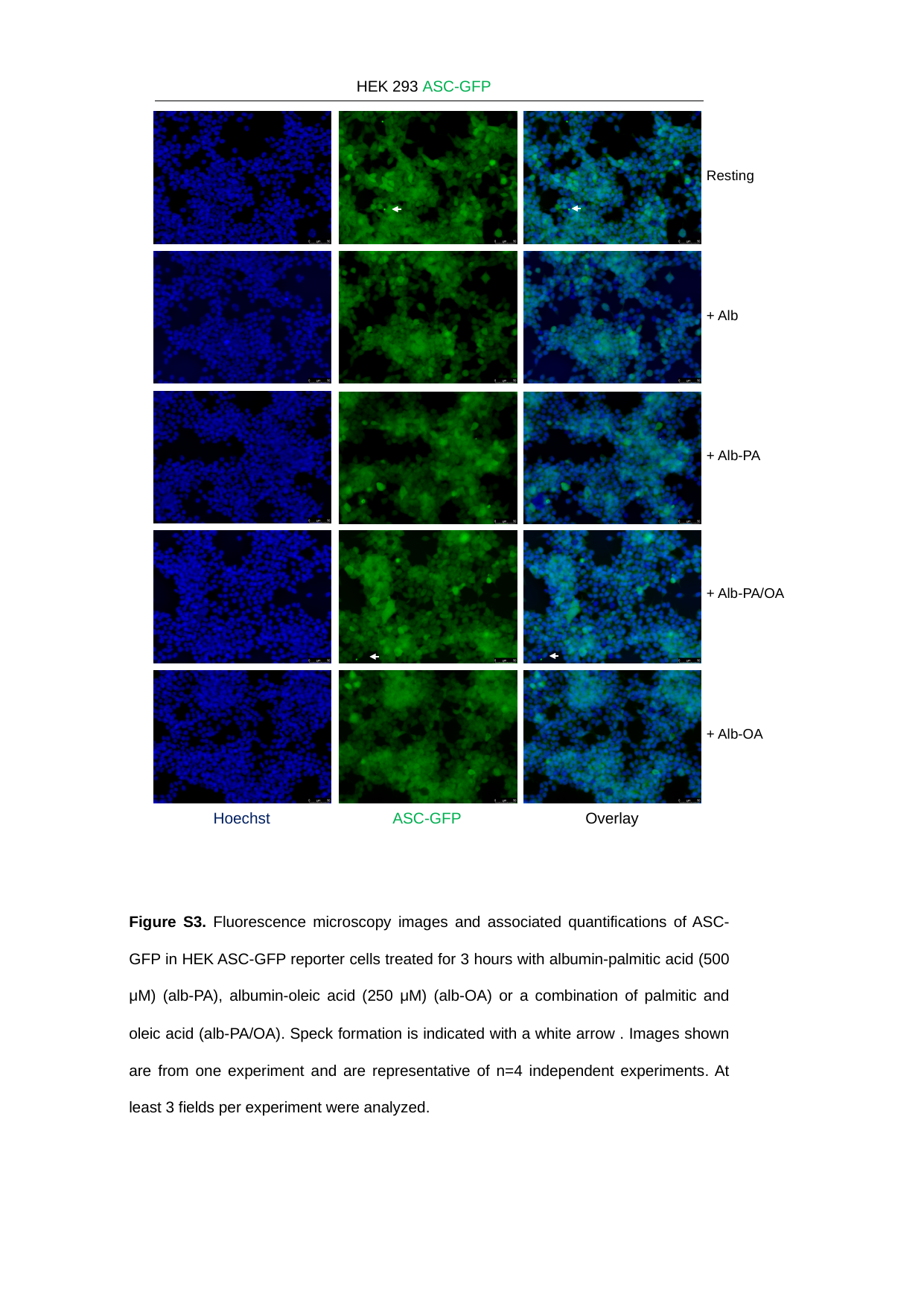

HEK 293 ASC-GFP
Resting
+ Alb
+ Alb-PA
+ Alb-PA/OA
+ Alb-OA
ASC-GFP
Overlay
Hoechst
Figure S3. Fluorescence microscopy images and associated quantifications of ASC-GFP in HEK ASC-GFP reporter cells treated for 3 hours with albumin-palmitic acid (500 μM) (alb-PA), albumin-oleic acid (250 μM) (alb-OA) or a combination of palmitic and oleic acid (alb-PA/OA). Speck formation is indicated with a white arrow . Images shown are from one experiment and are representative of n=4 independent experiments. At least 3 fields per experiment were analyzed.
