## Supplementary tables for "NLRP1 is activated by palmitic acid and induced in human metabolic dysfunction-associated steatohepatitis"

**Supplementary Table S1.** Primer sequences

| Gene Name | Gene symbol^a^ | Forward | Reverse |
| --- | --- | --- | --- |
| NLR family pyrin domain containing 1 | *NLRP1* | CACAGAAATCAGAGAAAGAGAG | AAATCCTCATTTTTCCAGGG |
| NLR family pyrin domain containing 3 | *NLRP3* | GATCTTCGCTGCGATCAACAG | CGTGCATTATCTGAACCCCAC |
| Tumor necrosis factor | *TNF* | GAGGCCAAGCCCTGGTATG | CGGGCCGATTGATCTCAGC |
| Interleukin 6 | *IL6* | ACTCACCTCTTCAGAACGAATTG | CCATCTTTGGAAGGTTCAGGTTG |
| Interleukin 1 beta | *IL1B* | ATGATGGCTTATTACAGTGGCAA | GTCGGAGATTCGTAGCTGGA |
| RNA, 18S ribosomal | *RNA18S1* | CCTCCAATGGATCCTCGTTA | AAACGGCTACCACATCCAAG |

^a^: HGNC, Hugo Gene Nomenclature Committee.

**Supplementary Table S2.** Characteristics of the study population

|  | No MASLD  (n=53) | MASL  (n=43) | MASH  (n=54) | *p value* |
| --- | --- | --- | --- | --- |
| Gender (F/M) | 50/3 | 29/14 | 34/20 | **<0.001** |
| Age (years) | 43.11 ± 10.34^a^ | 46.95 ± 11.39^a,b^ | 48.37 ± 10.22^b^ | **0.033** |
| Weight (kg) | 117.11 ± 18.84 | 121.39 ± 21.16 | 119.41 ± 24.74 | 0.685 |
| BMI (kg/m^2^) | 43.27 ± 5.69 | 44.21 ± 5.33 | 43.42 ± 6.77 | 0.522 |
| WC (cm) | 122.42 ± 12.98^a^ | 128.18 ± 12.91^b^ | 127.90 ± 15.23^a,b^ | **0.032** |
| SBP (mm Hg) | 132.73 ± 17.43^a^ | 142.14 ± 21.30^b^ | 141.43 ± 19.23^b^ | **0.010** |
| DBP (mm Hg) | 84.85 ± 11.71 | 85.93 ± 12.69 | 84.74 ± 12.06 | 0.668 |
| Glucose (mg/dL) | 88.02 ± 10.74^a^ | 99.84 ± 24.47^b^ | 115.65 ± 46.69^b^ | **<0.001** |
| HbA1c (%) | 5.58 ± 0.71^a^ | 6.03 ± 1.07^a,b^ | 6.36 ± 1.47^b^ | **<0.001** |
| Insulin (μIU/mL) | 13.38 ± 11.24^a^ | 13.87 ±6.71^a,b^ | 21.04 ± 21.07^b^ | **0.018** |
| HOMA-IR | 2.98 ± 2.60^a^ | 3.45 ± 1.99^a,b^ | 6.08 ± 6.91^b^ | **<0.001** |
| Cholesterol (mg/dL) | 159.92 ± 31.29 | 164.58 ± 33.00 | 163.26 ± 31.59 | 0.757 |
| HDL-c (mg/dL) | 46.90 ± 10.62^a^ | 38.93 ± 8.78^b^ | 40.46 ± 11.64^b^ | **<0.001** |
| LDL-c (mg/dL) | 87.92 ± 28.44 | 86.73 ± 29.18 | 85.10 ± 31.24 | 0.894 |
| Triglycerides (mg/dL) | 166.37 ± 125.88^a^ | 224.37 ± 192.63^b^ | 196.65 ± 90.59^b^ | **<0.001** |
| AST (U/L) | 16.96 ± 5.47^a^ | 19.20 ± 7.04^a^ | 26.51 ± 15.82^b^ | **<0.001** |
| ALT (U/L) | 15.77 ± 7.07^a^ | 21.05 ± 10.49^b^ | 30.44 ± 14.67^c^ | **<0.001** |
| GGT (U/L) | 20.46 ± 16.55^a^ | 21.98 ± 19.38^a,b^ | 29.62 ± 26.63^b^ | **0.001** |
| ALP (U/L) | 73.77 ± 20.73 | 68.40 ± 22.24 | 72.45 ± 22.66 | 0.216 |
| Albumin (g/dL) | 3.96 ± 0.39^a^ | 4.04 ± 0.45^a,b^ | 4.19 ± 0.32b | **0.007** |
| Platelets (x10^3^/μL) | 265.58 ± 61.93 | 245.81 ± 59.69 | 269.49 ± 69.65 | 0.131 |

Values are presented as mean ± standard deviation (SD). *p values* were calculated using ANOVA test in those parameters with a normal distribution (i.e., age, cholesterol, LDL-c, and Albumin), considering *p* < 0.05 significant; whereas *p* values for parameters without a normal distribution were calculated using a Kruskal-Wallis test (considering *p* < 0.05 significant), followed by a Bonferroni post hoc analysis for the intergroup differences test in those parameters with a significant *p value*. ALP, alkaline phosphatase; ALT, alanine aminotransferase; AST, aspartate transaminase; BMI, body mass index; DBP, diastolic blood pressure; F, female; GGT, gamma-glutamyltransferase; HbA1c, glycated hemoglobin; HDL-c, high-density lipoprotein cholesterol; HOMA-IR, homeostatic model assessment of insulin resistance; LDL-c, low-density lipoprotein cholesterol; M, male; MASH, metabolic dysfunction-associated steatohepatitis; MASL, metabolic dysfunction-associated steatotic liver; MASLD, metabolic dysfunction-associated steatotic liver disease; SBP, systolic blood pressure; WC, waist circumference. Different superscript letters indicate statistically significant difference at *P* < 0.05 within each row between the groups.

**Supplementary Table S3.** Histopathological findings in livers of the study population.

|  | MASL  (n = 43) | MASH  (n = 54) | *P value* |
| --- | --- | --- | --- |
| SAF score (0/1/2/3/4/5/6/7/8) | (0/19/16/7/1/0/0/0/0) | (0/0/0/20/13/11/6/2/2) | ˂ 0.001 |
| Steatosis (0/1/2/3) | (0/32/11/0) | (0/30/13/11) | 0.006 |
| Hepatocellular ballooning (0/1/2) | (25/17/1) | (0/42/12) | ˂ 0.001 |
| Lobular inflammation (0/1/2/3) | (40/3/0/0) | (0/42/12/0) | ˂ 0.001 |
| Fibrosis (0/1/2) | (43/0/0) | (42/11/1) | 0.004 |

Values are presented as frequencies. *P*-values were calculated using Pearson’s Chi-squared test considering *p* < 0.05 as significant. SAF, Steatosis, Activity and Fibrosis; MASL, metabolic dysfunction-associated liver, MASH, metabolic dysfunction-associated steatohepatitis.
